# Evolution of giant pandoravirus from small icosahedral viruses revealed by CRISPR/Cas9

**DOI:** 10.1101/2022.08.18.504477

**Authors:** Hugo Bisio, Matthieu Legendre, Claire Giry, Nadege Philippe, Jean-Marie Alempic, Sandra Jeudy, Chantal Abergel

**Author notes:** These authors contributed equally (Hugo Bisio, Matthieu Legendre). Correspondence (H.B.), (C.A.).

## Abstract

Giant viruses (GVs) are a hotspot of unresolved controversies since their discovery, including the definition of “Virus” and the existence of a fourth domain of life^1-3^. While increasing knowledge of genome diversity has accumulated^4^, functional genomics was largely neglected. Here, we describe an experimental framework to genetically modify nuclear GVs and its host *Acanthamoeba castellanii* using CRISPR/Cas9, allowing us to uncover the evolution from small icosahedral viruses to amphora-shaped GVs. Ablation of the icosahedral major capsid protein in the evolutionary intermediate mollivirus highlights a stepwise transition in virion shape and size. We additionally demonstrate the existence of a reduced core essential genome in pandoravirus, reminiscent of their proposed smaller ancestors. Genetic expansion led to increased genome robustness, indicating selective pressures for adaptation to uncertain environments. Overall, we introduce new tools for manipulation of the unexplored genome of nuclear GVs and demonstrate that viral gigantism can arise as an emerging trait.

---

Giant viruses (GV) have genomes up to 2.5 megabases and form viral particles that can match in size some cellular organisms^5,6^. Their dsDNA genomes can encode more than 1000 genes, 70% of them corresponding to proteins unseen in any other organisms^6,7^. These genes are referred to as ORFans^8^. In addition, these atypical viruses encode genes involved in translation and metabolism, hallmark functions of the cellular world^3,9,10^. These unprecedented features have led to great controversies regarding the origin of such viruses and blurred the division between cellular living organisms and viruses^1^. The origin of these viruses has been attributed either to the reduction evolution branching from the cellular world^1^ or the evolution towards complexity from smaller viruses^11^.

To achieve genetic manipulation of GVs, we built an *Acanthamoeba castellani* CRISPR/Cas9 system, using a plasmid encoding a polycistronic tRNA-gRNA^12,13^ and the *Sp*Cas9 fused to green fluorescent protein (GFP)^14^ (Fig. 1a). To test the system, we conducted Cas9-mediated modification of an episomally encoded red fluorescent protein (mRFP) (Fig. S1a). When amoebas were transfected with Cas9 and specific guides against the mRFP (Fig. S1a), we observed a significant decrease in the mRFP positive population, which correlated with Cas9 expression (GFP positive) (Fig. 1b-c). Double labeled amoebas were not observed when on-target guides were present (Fig. 1b-c). Accordingly, genotyping confirmed gene locus modification in ∼50% of the screened library (Fig. 1d and S1b). In order to increase the efficiency of genome modification, we hypothesized that the introduction of a “mutagenic chain reaction” (Fig. S1c)^15^ would allow the propagation of recombinant genes. Concordantly, 100% reduction of mRFP expression was demonstrated when this strategy was utilized (Fig. 1b-c and S1d). Correct integration and homozygosis were showed by PCR (Fig. 1e). No recombinant cells were detected when off-target guides were transfected (Fig. 1b-c), indicating that homologous recombination (HR) is rather inefficient in absence of double strand breaks.

**Figure 1.**
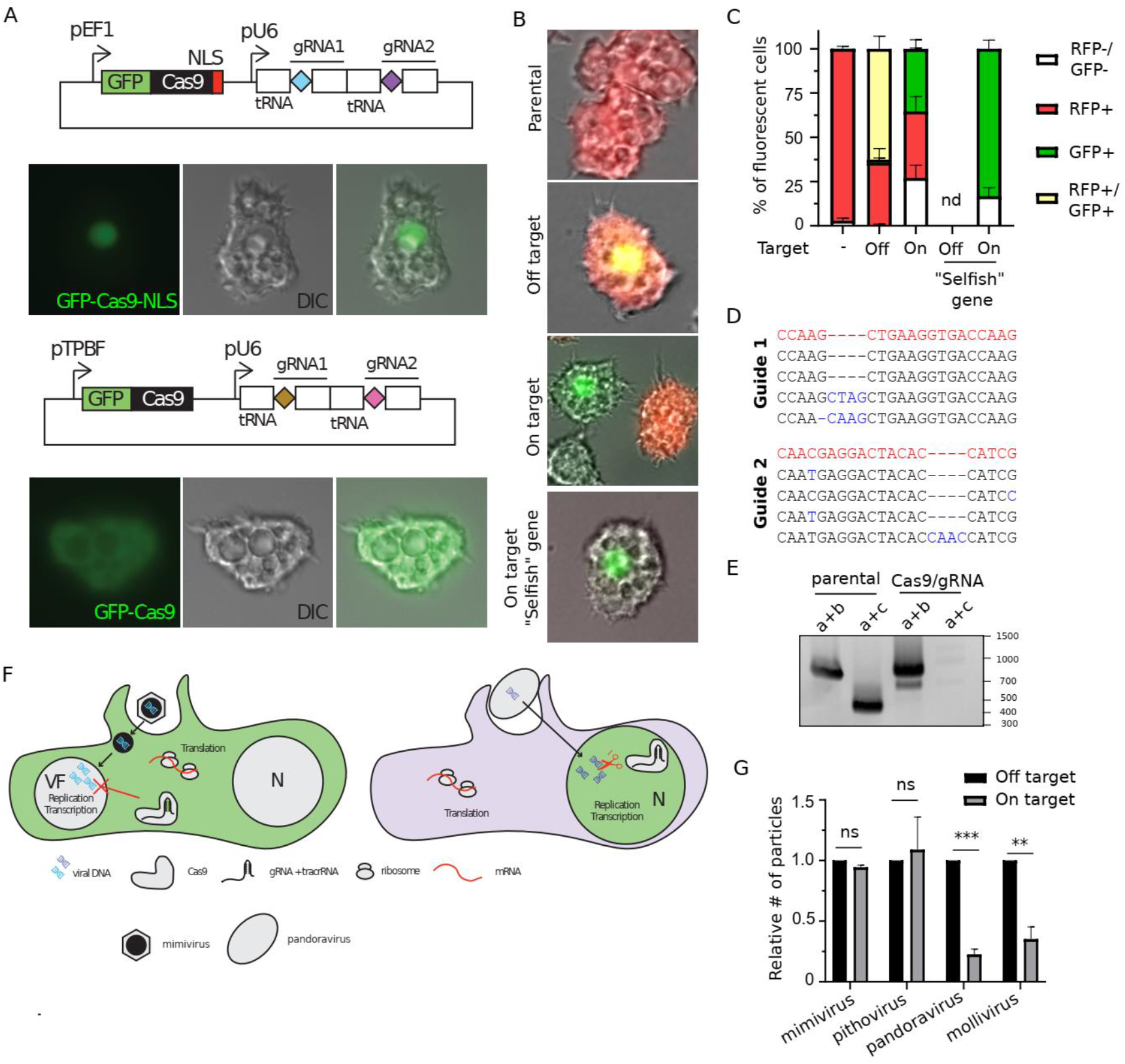
CRISPR/Cas9 allows manipulation of host and nuclear GVs. (a).Constructs used to constitutively express Cas9 in *A. castellanii*. The sequence of the gRNA is depicted with diamonds followed by the Cas9-binding scaffold and an *A. castellanii* tRNA (white rectangles). The localization of the GFP-Cas9 fusion for the different constructs is shown by fluorescence. (b).Representative micrographs showing amoebas after selection for 2-3 weeks with the appropriate drug(s) (refer to materials and method). mRFP (product of the targeted gene) and GFP (Indicating Cas9 expression) fluorescence are shown. (c).The quantification of the micrograph shown in B. The mean ± SD (3 independent transfections) of non-fluorescent, mRFP+, GFP+ and GFP+mRFP+ cells is shown. (d).Representative sequencing results of targeted guide sequences on *rfp* upon transfection with on-target gRNAs. PCR were performed on the target sequences as shown in Figure S1a-b, cloned into a TA cloning vector and single clones were sent for sequencing. The wild type sequence and mutations generated are shown in red and blue respectively. (e).Gene disruption of *rfp* by the “mutagenic chain reaction” observed at the population level after 2-3 weeks post transfection. Expected PCR size: a+b: 837bp (wt), a+b 890bp (KO), a+c: 500bp (wt). A map of the construct used and a cartoon depicting the strategy of disruption is shown in Figure S1c-d. (f).Schematic depiction of genetic manipulation of GVs. Recombinant *A. castellanii* that express a nuclear or cytoplasmic Cas9 are shown in the left and right, respectively. CRISPR/Cas9 allows manipulation of nuclear GVs but not cytoplasmic GVs. (g).Relative quantification of the number of viruses produced 24 hours post infection (hpi) of CRISPR/Cas9 expressing amoebas. A cytoplasmic version of Cas9 was used to attempt gene disruption of cytoplasmic GV (mimivirus reunion^19^ and pithovirus sibericum^18^) and a nuclear version of Cas9 was used for the gene disruption of nuclear GVs (pandoravirus neocaledonia^7^ and mollivirus Kamchatka^17^). Guides off-target and on-target of the *rbp1* gene for each virus were used. Data correspondion to the mean ± SD of 3 independent experiments. p-value: <0.05 (*), <0.01 (**), <0.001 (***) and <0.0001 (****).

All giant viruses encode an RNA polymerase (RNAP), which modeled the RNAPII of modern eukaryotes^16^. Due to its likely essentiality for viral replication, we targeted the *rpb1* gene to assess the capacity of Cas9 to target viral genomes (Fig. S1e). Nuclear Cas9 expression in amoeba cells (Fig. 1a) showed efficient regulation of pandoravirus neocaledonia^7^ and mollivirus kamchatka^17^ replication (Fig. 1f-g), while the cytosolic version of the protein (Fig. 1a) was unable to target the genome of pithovirus sibericum^18^ or mimivirus reunion^19^ (Fig. 1f-g). The incapability of Cas9 to target the genome of strictly cytoplasmic viruses might reside in the shielding of the DNA by these viruses, excluding GFP-Cas9 from the viral factory as shown by fluorescence (Fig. S1f). A similar strategy was previously demonstrated for some bacteriophages^20^. Coherently with the likely essentiality of the *rnap*, only virions containing wild-type genomes (which escaped Cas9 targeting) were recovered in genomic libraries (Fig. S1g-h).

To gather information on the repair mechanisms of double strand breaks in nuclear GVs, we targeted genes with suspected different degrees of importance for virus replication. Fitness reduction of pandoravirus was highly dependent on the targeted gene locus (Fig. 2a). Surprisingly, targeting of genes at the 3’ of the genome resulted in large deletions rather than discrete modification of the targeted locus (Fig. 2b-d and S2a), suggesting these viruses are not efficient in repairing double strand breaks.. Quantification of the ratio between PCR products using 5’ and 3’ regions of the genome as templates demonstrates that large deletions are rather frequent at the 3’ of the genome, but not detectable at the 5’ (Fig. 2b). Moreover, viruses harboring such 3’ large deletions were able to complete the full cycle of replication since cloning was achievable (Fig. 2c). Importantly, pandoravirus was capable of losing at least over 600,000 bp (Δ1195458-1838505 base pairs) and more than 300 genes (Δpneo_612-974), as shown by pulsed field gel electrophoresis (Fig. 2d) and PCR amplification of specific loci (Fig. S2a). Interestingly, while deletion of almost one third of the genome led to a significant reduction on viral fitness (Fig. 2e), no changes in virion morphology were observed (Fig. 2f). This data strongly correlates with the strikingly low abundance of virion proteins encoded by genes located at the 3’
s end of the genome (Fig. 2g). Moreover, viral DNA replication was significantly impaired in viruses lacking the genes present at the 3′ of the genome (Fig. 2h). Taken together, this data demonstrates that these genes are important during the intracellular stage of the infectious cycle, most likely for host pathogen interaction.

**Figure 2.**
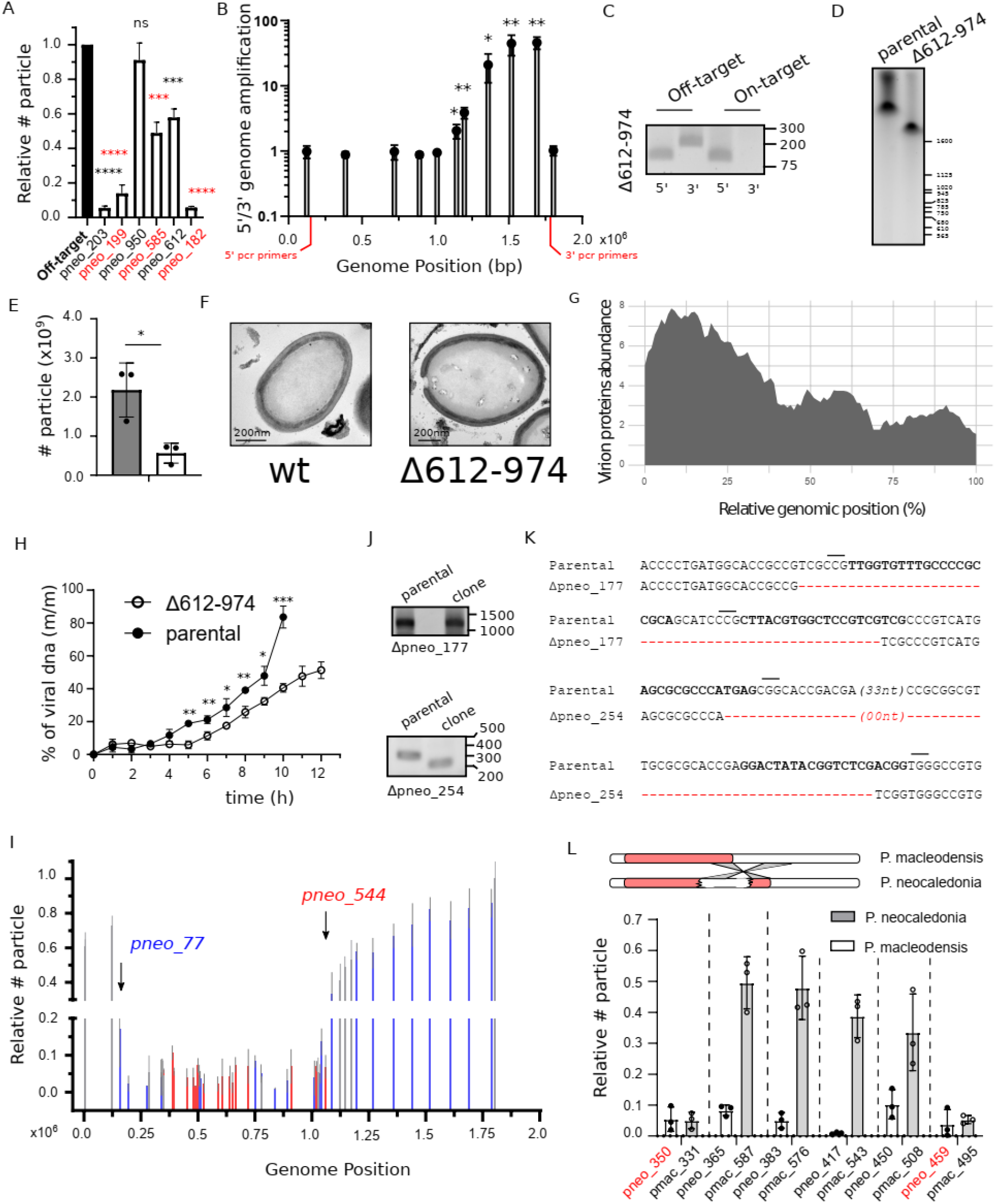
CRISPR/Cas9 targeting of pandoravirus genome demonstrates the presence of a core essential genome. (a).Relative quantification of the number of viruses produced 24 hpi of CRISPR/Cas9 expressing amoebas. Guides off-target and on-target of different genes were used. Data correspond to the mean ± SD of 3 independent experiments. (b).qPCR analysis indicates the presence of virus with large deletions at the 3’ of the genome. A ratio of the copy number of genes pneo_74 (5’) and pneo_974 (3’) is shown and their location are marked in red. An intact genome corresponds to the one with a ratio close to 1. The location of the bars indicates the position of the CRISPR/Cas9 targeted gene on the genome of Pandoravirus neocaledonia. Data correspond to the mean ± SD of 3 independent experiments. (c).Clonality of viral strains with large deletions at the 3’ of the genome was analyzed by PCR. Expected PCR size: 5’: 150bp (*pneo_74*), 3’: 220bp (*pneo_974*). Genetic loss of other target genes is show in Figure S2D. (d).Pulsed field gel electrophoresis of the genomic DNA of pandoravirus neocaledonia. Genome obtained from parental virions (2.05 Mbp) or virions containing a large deletion at the 3’ end of the genome (Δ612-974) are shown. (e).Quantification of the number of viruses produced 24 hpi. Data correspond to the mean ± SD of 3 independent experiments. Parental strain is shown in gray while mutant lacking genes 612 to 974 are represented in white. (f).TEM image of an ultrathin section of p. neocaledonia viral particle. (g).Abundance of proteins detected in the p. neocaledonia virion proteome according to the genomic location the cognate genes. Protein abundance was computed from the normalized log2(iBAQ) values from ^7^ and averaged over the genome using a sliding window. (h).DNA replication was analyzed by qPCR. Viral DNA is represented as a percentage of total DNA in the sample (mass of viral DNA versus total DNA mass (m/m)). Data correspond to the mean ± SD of 3 independent experiments. (i).Relative quantification of the number of viruses produced 24 hpi of CRISPR/Cas9 expressing amoebas ranked by their position in the genome. Guides off-target and on-target of different genes were used. Data correspond to the mean ± SD of 3 independent experiments. Genes confirmed to be essential, dispensable or not analyzed are represented in red, blue and gray, respectively. The approximate limits of the essential core of the genome are marked with arrows and the gene targeted indicated. (j).Gene disruption of *goi* is demonstrated by PCR. Expected PCR size: pneo_177: wt, 1250bp – mutant, 1200bp; pneo_254: wt, 320bp – mutant, 220bp. Cartoon depicting the strategy for genotyping is shown in Figure S3A. (k).Sequencing results of targeted region on clonal populations of Δ*pneo_177* and Δ*pneo_254*. PCR were performed on the target sequences as shown in Figure 3D. The deleted sequences are shown in red. gRNA sequences are highlighted in bold and PAM sequences marked with a line. (l).Relative quantification of the number of p. neocaledonia and p. macleodensis produced 24 hpi of CRISPR/Cas9 expressing amoebas ranked by their position in the genome of p. neocaledonia. Essential genes in p. neocaledonia are highlighted in red. Orthologous genes between p. neocaledonia and p. macleodensis are shown in pairs. A schematic representation of the genome organization of p. neocaledonia and p. macleodensis is also shown. The limits of the inverted region of the genome is shown in gray. The essential core of the genome is represented with a red box. p-value: <0.05 (*), <0.01 (**), <0.001 (***) and <0.0001 (****).

Interestingly, fitness differences during the first generation of viral production upon targeting with CRISPR/Cas9 were associated to the gene location rather than the gene target itself (Fig. 2i). This data indicates that large deletions were not viable when generated at the 5’ of genome (core genome). The core genome includes the genes pneo_77 (dispensable in blue, Fig. 2i) to pneo_544 (essential in red, Fig. 2i). The identification of the precise limits of this area would require further characterization. Meanwhile, amplifications of the viruses produced under a constant selection with Cas9 by targeting the core genome led occasionally to the production of resistant viruses (Fig. S2b) which contain modification of the genome at the target site (Fig. 2j). Coherently, shorter deletions were observed in viruses when genes present at the core genome were targeted (Fig. 2k), demonstrating the capability of these viruses to repair by non-homologous end joining, even if not efficiently. Taken together, we were able to determine a core essential region of the genome associated to the 5’ (with a high density of essential genes, labeled in red), and a non-essential one located at the 3’ of the genome (labeled in blue) (Fig. 2i). Importantly, not all genes present at the core genome are essential since knock out of some of these genes was accomplished (Fig. 2i-k). This essential core region was even smaller in p. macleodensis, which lacks a recent genomic inversion present in p. neocaledonia (Fig. 2l). This inversion is located at the intersection of the essential and non-essential zone of the genome (Fig. 2i)^7^. Concordantly, this inversion in p. neocaledonia has introduced a fragment of the non-essential region of the genome into a newly fragmented essential core region of its genome (Fig. 2l).

Importantly, the assessment of the non-essential region showed an increase fitness cost upon larger deletions, as shown by a larger amount of progeny produced when the targeting site is present closer to the 3’ end of the genome. As a result, this data indicates a cumulative effect of the loss of function of the genes contained in the non-essential region of the genome (Fig. 2i).

The distribution of essential genes observed in pandoravirus would be parsimoniously explained by the expanding genome hypothesis and rather difficultly with the reductive one. Concordantly, reductive evolution in pathogenic bacteria or eukaryotes did not lead to the concentration of essential genes in particular regions of their genomes^21,22^. In order to test the genome expansion hypothesis in pandoravirus, we classified genes according to their conservation in their distant relatives of the *Nucleocytoviricota* (Fig. 3a)^11,16^, and analyzed their position in the genome of p. neocaledonia (Fig. 3b). p. neocaledonia genes with orthologs in *Phycodnaviridae* and mollivirus are enriched at the 5’ of the genome, coinciding with the essential core of the genome (Fig. 3b). On the other hand, clade-specific genes are more uniformly distributed and strain-specific genes (ORFan genes) tend to accumulate at the 3’ end of the genome (Fig. 3b). Concordantly, this data supports the hypothesis of a biased expansion of the pandoravirus genome from a *Phycodnaviridae*-like virus towards the 3’ end of the genome.

**Figure 3.**
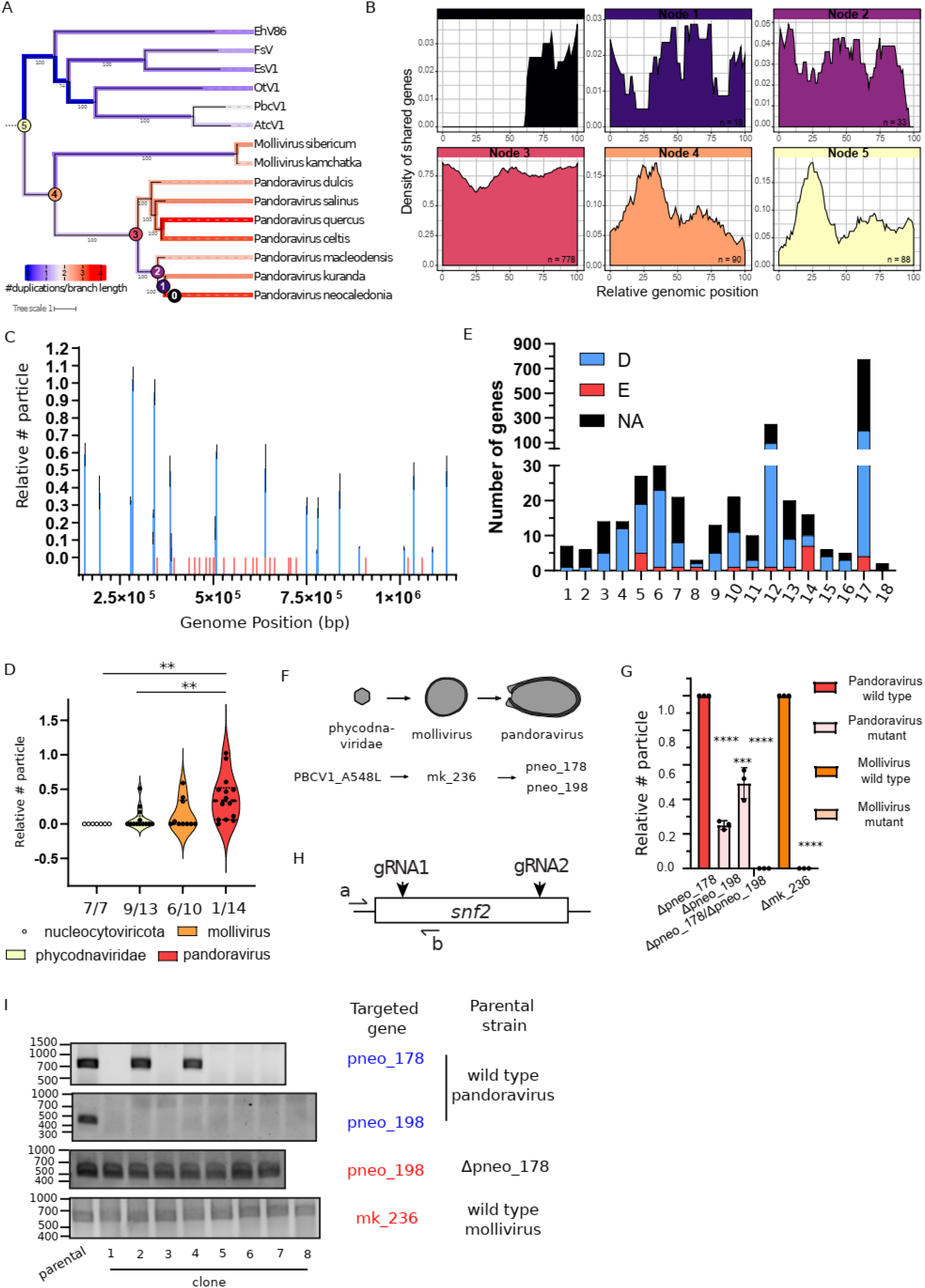
Pandoravirus genome organization keeps traces of their smaller ancestors. (a.Phylogenetic tree of Phycodnaviridae, Molliviridae and Pandoraviridae computed from a concatenated multiple alignment of 435 orthologous proteins present in at least two viruses.Bootstrap values were computed using the ultrafast boostrap method^48^. Number of gene duplication events along the branches from the phylogenetic tree are also shown. The numbers were normalized by branch length, log10 transformed and color-coded from blue (low number of duplications) to red (high). (b).Density of P. Neocaledonia genes along the genome according to its predicted ancestry. The P. neocaledonia genes were classified based on the presence of homologs in other viruses and color-coded from the corresponding ancestral node shown in panel A. (c).Relative quantification of the number of viruses produced upon knock-out of genes located at the essential core of the genome, ranked by their position in the genome. Data correspond to the mean ± SD of 3 independent experiments. Genes that were knocked out are highlighted in blue, while genes that could not be knocked out are shown in red. (d).Correlation between the number of particles produced 24 hpi upon single gene knock out and proposed depth of conservation. The phylogenetic relationship of p. neocaledonia with other nucleocytoviricota is illustrated in Figure 3a. The distribution of phenotype scores in each category is plotted and the number of genes analyzed for each category is showed on the bottom of each violin plot. Bars indicate the group median. The number of essential genes over the total of genes analyzed in each group is also indicated. (e).Correlation between dispensability/essentiality (D = dispensable, E = essential, NA = not analyzed) and functional annotation of pandoravirus neocaledonia genes. Gene ontology classes: 1-Carbohydrate metabolism. 2-Cofactor biosynthesis. 3-Collagen. 4-Cytoskeleton. 5-DNA replication, repair, packaging and modification. 6-DUF. 7-Kinases/phosphatases, ubiquitin (post-translational modifications). 8-Lipids binding. 9-Misc enzymes. 10-Nucleotide metabolism and binding. 11-Proteases and protease inhibitor. 12-Protein-protein interactions (Ankyrin, F-box, Morn, BTB-Poz, Pan Apple). 13-Redox and metallo enzymes. 14-RNA transcription and regulation. 15-Translation and amino-acid metabolism. 16-Transport. 17-Uncharacterized. 18-Virulence factors. (f).Transition from mollivirus to pandoravirus coincides with the duplication of a SNF2-like helicase. (g).Relative quantification of the number of viruses produced upon knock-out of *snf2* like genes in pandoravirus or mollivirus. Data correspond to the mean ± SD of 3 independent experiments. (h).Cartoon depicting the strategy for genotyping shown in (Fig. 3i). (i).Gene presence or disruption of *goi* is demonstrated by PCR in different clones obtained upon gene targeting by Cas9. PCR positive clones represent viruses which escape Cas9 targeting. Expected pcr size: pneo_178: 790bp; pneo_198: 466bp; mk_236: 562bp. p-value: <0.05 (*), <0.01 (**), <0.001 (***) and <0.0001 (****).

Importantly, gene persistence throughout evolution correlates with the fitness cost associated to the loss of function of that gene^23^. Thus, to further test the genome expansion theory of pandoravirus, we attempted gene KO for approximately 10% of the genes present in the essential region of the genome (Fig. 3c and S3). In agreement with the genome expansion theory, the density of essential genes increased at the positions where *Phycodnaviridae* and mollivirus conserved genes are also highly represented (Fig. 3c). Moreover, the assessment of the depth of conservation of genes across *Nucleocytoviricota* versus the phenotype associated to their deletion indicates that shared genes have a greater tendency to produce a more drastic effect upon deletion (Fig. 3d). This distribution of phenotypes followed the trend predicted by the genome expansion theory and correlates with the proposed evolutionary pathway from a phycodnavirus-like ancestor to pandoravirus (Fig. 3a). Moreover, genes largely conserved across the *Nucleocytoviricota* (despite multiple independent deletions rendered only one gene strictly conserved in all viruses^24^) and used for the phylogenetic reconstruction of the phylum are essential for pandoravirus replication (Fig. 3d). These genes include: B DNA Polymerase (PolB), A32-like packaging ATPase (A32), virus late transcription factor 3 (VLTF3), large RNA polymerase subunits (RNAP1 and RNAP2), TFIIB transcriptional factor (TFIIB) and D5-like primase-helicase^11,16,24^. This data aligns with their use for phylogenetic reconstructions by indicating their likely ancestry for the phylum.

Finally, analysis of gene essentiality according to gene broad functional annotations indicates that essential genes accumulate particularly under “replication” and “transcription” functions, while “Translation and amino-acid metabolism” includes none (Fig. 3e). This observation further supports the viral origin of pandoravirus and indicates that translation-related genes might have been acquired recently to maximize expression of viral genes. Genes with unknown functions, including all the ORFan genes, englobe a large amount of non-essential genes. Thus, these genes are also likely to be recent acquisitions of the pandoravirus.

Previous reports have shown an unexpected prevalence of multiple-copy genes in Pandoravirus genomes^7^. Moreover, the speed of acquisition of these multi-copy genes increased with the genome expansion (Fig. 3a). This data suggests that genome expansion was accompanied by a strong increase in genetic redundancy and thus, led to increased robustness of the biological system^25^. In order to test this hypothesis, we analyzed a family of SNF2-like genes present in phycodnaviridae, mollivirus and pandoravirus (Fig. 3f). SNF2 genes are helicase-related ATPases which in eukaryotic cells drive chromatin-remodeling ^26^. While *Phycodnaviridae* and mollivirus encode only one SNF2-like gene, this gene is present in two copies in the genome of pandoravirus (Fig. 3f). Interestingly, phylogenetic analysis of these genes indicates that the duplication occurred in the ancestor of pandoravirus and mollivirus (Figure S4). A followed-up deletion of one of these genes would explain the presence of a single gene in mollivirus (Figure S4). Such evolutionary history can be explained by the proposed “genomic accordion” evolution proposed for GVs^27,28^. Importantly, both genes present in pandoravirus were shown dispensable individually (Fig. 3g) and viruses encoding a deletion of these genes easily cloned (Fig. 3h-i). On the other hand, double KO led to synthetic lethality (Fig. 3g-i). Concordantly, this data demonstrates the presence of epistatic effects and genetic redundancy in GVs. In order to assess if genetic redundancy increased with genome expansion, we targeted the SNF2-like gene in mollivirus, which is present in a single copy (Fig. 3f). We were was unable to be knocked out *mk_236*, which strongly supports its essentiality (Fig. 3g-i). Thus, such redundancy displayed by pandoravirus would indicate that the flexibility for adaptation to uncertain environments might generate a strong selective pressure in these viruses.Concordantly, an increased genetic robustness might be a key component which explains the selective advantage that led to viral giantism.

Viruses belonging to the *Nucleocytoviricota* tended to lose some of the core genes independently throughout evolution^11,24^. In particular, pandoravirus lacks a Nucleocytoviricota signature MCP protein^11^. MCP is a major component of the viral particles of icosahedral viruses and key factor for the shaping of their capsids^29^. Interestingly, despite its spherical shape, mollivirus encodes an MCP-like protein^30^, and it has been speculated that this gene would be a trace of their icosahedral ancestry (Fig. 4a)^31^. AlphaFold predictions^32^ confirmed the jelly-roll folding unit of *ms_334* which closely resembles the MCP of other icosahedral viruses like Paramecium bursaria chlorella virus 1 (PBCV-1)^29^ (Fig. 4b). To assess the role of this protein in the formation of mollivirus particles, we first expressed an ectopic copy of the gene in the amoeba. Fluorescent N- or C-terminally tagged MCP localized into the cytoplasm of non-infected amoebas but re-localized to specific sub-compartments upon infection by mollivirus (Fig. 4c). Moreover, incorporation of MCP to the viral particle was confirmed by fluorescence (Fig. 4d). In order to attempt genetic deletion of mollivirus MCP, we infected transgenic amoebas expressing Cas9 and gRNAs targeting the *mcp* gene in mollivirus. Targeting of *mcp* efficiently reduced viral fitness of mollivirus (Fig. 4e). Trans-complementation, as we have previously described^33^, was not successful to restore viral growth (Fig. 4f). Expression quantities or timing might explain the failure of trans-complementation. Thus, we developed a system for cis-complementation of gene KO (Fig. 4f). To achieve cis-complementation, we built a plasmid which encodes for a second copy of the *mcp* gene containing silent mutations at the gRNA target sites; and a nourseothricin N-acetyl transferase (NAT) resistance cassette, which confers resistance to nourseothricin^34^. The NAT selection cassette was expressed under the control of a pandoravirus promoter (Fig. 4f). Importantly, such recombinant viruses contain two copies of the *mcp*: one susceptible to cleavage by Cas9, and one resistant to its targeting (Fig. 4f and Fig. S5). Contrary to trans-complementation, cis-complementation efficiently restored viral fitness upon Cas9 cleavage of *mcp* (Fig. 4e). Moreover, while targeting of *mcp* did not impact DNA replication of the virus (Fig. 4g), biogenesis of virions was largely impaired (Fig. 4h). In contrast to untargeted or cis-complemented viruses, targeting of *mcp* on wild-type viruses led to the accumulation of membranous material in the cytoplasm and absence of mature neosynthesized virions (Fig. 4h). This data indicates a failure to initiate tegument biogenesis upon recruitment of an open cisterna^35^ and suggests a scaffolding function for mollivirus MCP. Overall, this data supports a stepwise replacement of the MCP functions during the evolution of phycodnavirus-like icosahedral viruses to mollivirus-like and later, pandoravirus. Which molecular changes in the particles of pandoravirus allowed the dispensability of the scaffolding functions of mollivirus MCP are yet to be identified.

**Figure 4.**
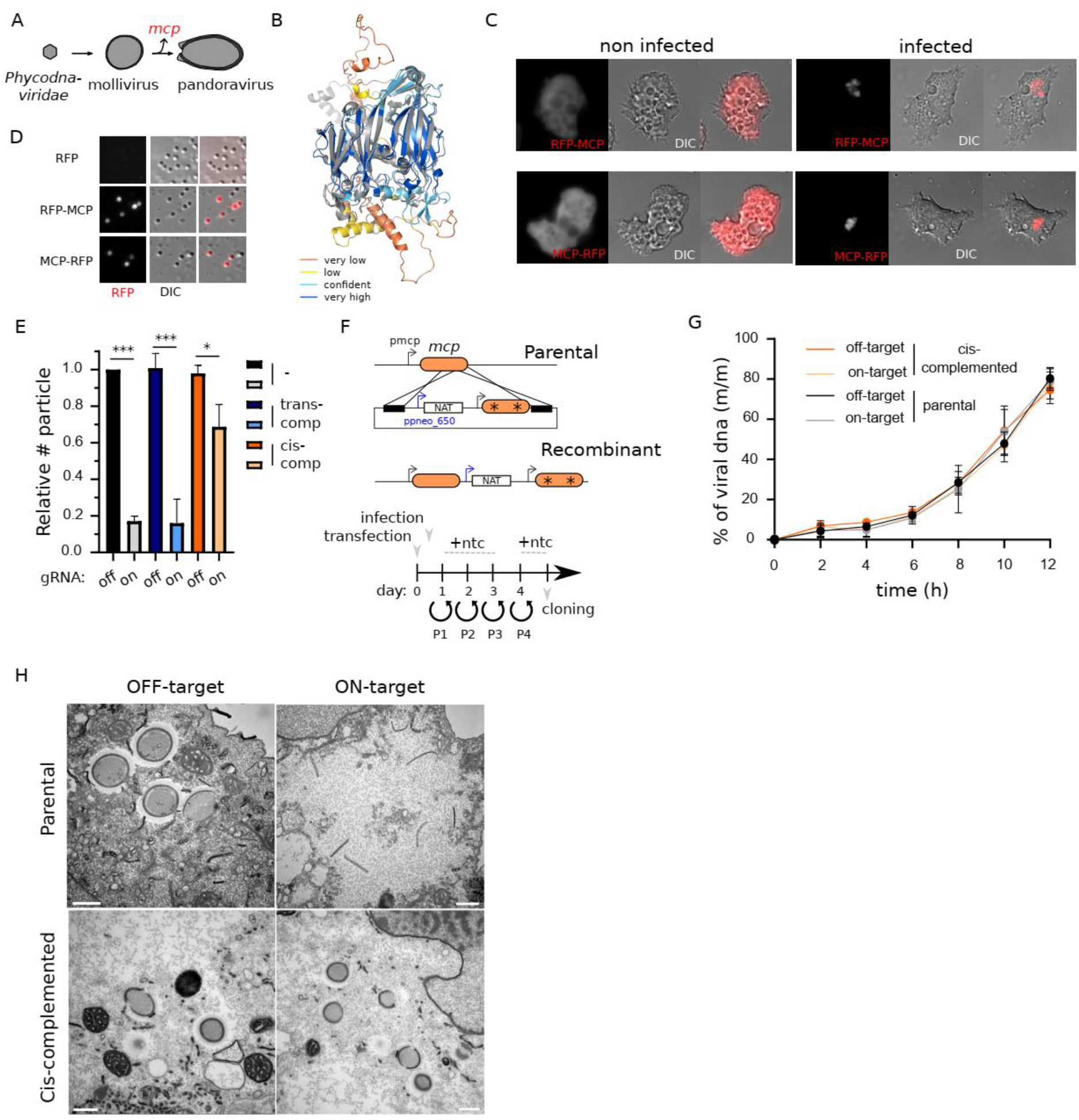
Mollivirus MCP acts as a scaffolding protein for tegument biosynthesis. (a).Schematic depiction of the proposed model of evolution for pandoravirus. Major capsid protein loss is indicated by a red color. (b).Alignment of the AlphaFold prediction of the mollivirus major capsid protein (mcp, ms_334) and the crystal structure of the PBCV-1 mcp (5TIQ). PBCV-1 mcp is shown in gray while mollivirus mcp is colored by its confidence score (pLDDT): Very low (0<pLDDT<50), low (50<pLDDT<70), confident (70<pLDDT<90) and high (90<pLDDT<100). (c).Light fluorescence microscopy images of *A. castellanii* cells expressing N- or C-terminally RFP tagged MCP. Representative images of non-infected cells and cells infected with mollivirus are shown. Images were obtained 6hpi. (d).Light fluorescence microscopy images of mollivirus produced in *A. castellanii* cells expressing RFP or N- or C-terminally RFP tagged MCP. (e).Relative quantification of the number of viruses produced upon knock-out of *mcp* gene in mollivirus. Cis-complementation was accomplished as shown in Figure 4F. Data correspond to the mean ± SD of 3 independent experiments. * indicates the presence of silent mutations on the sequence. (f).Schematic representation of the vector and strategy utilized for *mcp* KO cis-complementation. Selection cassette was introduced by homologous recombination and recombinant viruses were generated, selected and cloned as indicated by the timeline. Viral infection was performed 1 hour post-transfection. Ntc: Nourseothricin. * represents the presence of silent mutations at the gRNA targeting site. (g).DNA replication was analyzed by qPCR. Viral DNA is represented as a percentage of total DNA in the sample. Data correspond to the mean ± SD of 3 independent experiments. (h).Electron microscopy imaging of the mollivirus replication cycle in *A. castellanii*. Images were acquired 8hpi and mollivirus were used to infect cells expressing Cas9 and gRNA on- or off-target of the *mcp*. Parental or cis-complemented viral strains were used for infection in parallel experiments. p-value: <0.05 (*), <0.01 (**), <0.001 (***) and <0.0001 (****).

The study of the origin of viruses and their evolutionary history remains rather challenging for a reliable phylogenetic reconstruction due to their fast evolution and quick exchange of genetic markers^11,36^. Regardless, such fast evolution in dsDNA viruses (with low mutation rates but high burst size^37^) might have been key to the evolution of all living cells^38,39^, and would have contributed to shape modern genes^16^, genetic elements^40^, or even organelles like the nucleus^41,42^. Here, we introduced a battery of tools which allows genetic manipulation of host and nuclear GV and thus, the trackability of these gene exchanges. This includes the characterization of the neglected functional genomic of nuclear GV and their up-to 70% ORFan gene-containing genome. ORFan genes have contributed substantially as evolutionary innovations^8^ and are expected to arise as lineage-specific adaptations to the environment^43,44^. The role of such innovations and unpredictable functions in pandoravirus are now open to rigorous investigation. Overall, our analysis demonstrates the potential of genetic screens to uncover conserved biological processes and host-pathogen interactions for the most genetically complex viruses discovered so far.

## Acknowledgments

We thank Jean-Michel Claverie for providing the samples to isolate Pandoravirus kuranda and critical discussion to improve the manuscript. This study was founded he European Research Council (ERC) under the European Union’s Horizon 2020 research and innovation program (grant agreement No 832601; N.P., S.J., and C.A.). H.B. is the recipient of an EMBO Long-Term Fellowship (ALTF 979-2019).

## Author contributions

H.B., conceptualization, methodology, validation, formal analysis, investigation, visualization, writing – original draft; M.L., conceptualization, methodology, software, validation, formal analysis, investigation, visualization, writing – review & visualization; C.G., investigation, writing – review & visualization; N.P., methodology, investigation, writing – review & visualization; J.M.A., methodology, investigation; S.J., methodology, investigation, writing – review & visualization; C.A., conceptualization, project administration, funding acquisition, writing – original draft.

## Materials and Methods

### Viral strains utilized in this work

The following viral strains have been used in this study: pandoravirus neocaledonia^7^, pandoravirus macleodensis^7^, pandoravirus kuranda (this study, ON887157), mollivirus kamchatka^17^, pithovirus sibericum^18^ and mimivirus *reunion*^19^.

### Cloning of DNA constructs

All primers used in this study are listed in Supplementary Table 2. All vectors used in this study are listed in Supplementary Table 3.

#### CRISPR/Cas9 expression vector

Vectors for CRISPR/Cas9 expression in *A. castellanii* cells were generated by inserting a codon optimized *Sp*Cas9 (with or without a nuclear localization signal (NLS)) and a cassette for polycistronic gRNA expression into the pEF1-GFP-NEO vector^33^. The cassette for polycistronic gRNA expression was synthesized by GenScript and designed to contain an *A. castellanii* U6 promoter-tRNA-gRNA1-tracerRNA1-tRNA-gRNA2-tracerRNA2. This cassette was further modified by PCR and circularized by recombination (InFusion Takara) to replace gRNA1-tracerRNA1-gRNA2 by a NotI site. gRNAs were included in primers synthesized by eurofins genomics, and inserted into the NotI site by InFusion (Takara).

#### MCP-mRFP and mRFP-MCP expression vectors

PCR product of the *mcp* was generated using *M. kamchatka* genomic DNA produced by Wizard genomic DNA purification kit (PROMEGA) according to manufacturer’s specifications. Primers used are HB84/HB85 and HB86/HB87. Vectors for expression of *mcp* were generated using the pEF1-mRFP-NAT vector by InFusion (Takara) into the NdeI or XhoI site for N- or C-terminal RFP tagging, respectively.

#### MCP second copy

The plasmid for cis-complementation was generated by sequential cloning of the 3’ UTR of pneo_480, the promoter of pneo_650, a Nourseothricin N-acetyl transferase (NAT) selection cassette, the 3’ UTR of pneo_650, the promoter of *mcp* and the *mcp* coding sequence. Each cloning step was performed using the Phusion Taq polymerase (ThermoFisher) and InFusion (Takara). Silent mutations at the gRNA targeting site were also introduced using InFusion. Finally, 500bp homology arms were introduced at the 5’ and 3’ end of the cassette in order to induce homologous recombination with the viral DNA. Prior to transfection, plasmids were digested with EcoRI and NotI.

### Cell culture and establishment of cell lines

#### Cell culture

*Acanthamoeba castellanii* (Douglas) Neff (American Type Culture Collection 30010TM) cells were cultured at 32 °C in 2% (wt/vol) proteose peptone, 0.1% yeast extract, 100μM glucose, 4mM MgSO_4_, 0.4mM CaCl_2_, 50 μM Fe(NH_4_)_2_(SO_4_)_2_, 2.5 mM Na_2_HPO_4_, 2.5 mM KH_2_PO_4_, pH 6.5 (home-made PPYG) medium supplemented with antibiotics [ampicilline 100 μg/mL, and Kanamycin 25 μg/mL]. 100 μg/mL Geneticin G418 or Nourseothricin was added when necessary.

#### Cell transfection

1.5×10^5^ *Acanthamoeba castellanii* cells were transfected with 6 μg of each plasmid using Polyfect (QIAGEN) in phosphate saline buffer (PBS). Selection of transformed cells was initially performed at 30 μg/mL Geneticin G418 or Nourseothricin and increased up to 100 μg/mL within a couple of weeks.

### Establishment of viral lines

#### Cas9 KO viral lines

Cell lines expressing Cas9 and gRNA were infected with a MOI (Multiplicity Of Infection) of 1. Infection was allowed to proceed for 24 hours. This new generation of viruses was used to quantify the number of particles produced or cloned for isolation of viral strains.

#### Insertion of MCP

1.5×10^5^ *Acanthamoeba castellanii* cells were transfected with 6 μg of linearized plasmid using Polyfect (QIAGEN) in phosphate saline buffer (PBS). One hour after transfection, PBS was replaced with PPYG and cells were infected with 1.5×10^7^ mollivirus particles for 1 hour with sequential washes to remove extracellular virions. 24h after infection the new generation of viruses (P0) was collected and used to infect new cells. An aliquot of P0 viruses was utilized for genotyping in order to confirm integration of the selection cassette. New infection was allowed to proceed for 1 hour, then washed to remove extracellular virions and nourseothricin was added to the media. Viral growth was allowed to proceed for 24 hours. This procedure was repeated one more time before removing the nourseothricin selection to allow viruses to expand more rapidly. Once, viral infection was visible, selection procedure was repeated one more time. Viruses produced after this new round of selection were used for genotyping and cloning.

#### Cloning and genotyping

150000 *A. castellanii* cells were seeded on 6 well plates with 2mL of PPYG. After adhesion, viruses were added to the well at a multiplicity of infection (MOI) of 2. One hour post infection, the well was washed 5 times with 1mL of PPYG and cells were recovered by well scraping. Amoebas were then diluted until obtaining a suspension of 1 amoeba/μL. 1μL of such suspension was added in each well of a 96-well plate containing 1000 uninfected *A. castellanii* cells and 200μL of PPYG. Wells were later monitored for cell death and 100μL collected for genotyping. Genotyping was performed using Terra PCR Direct Polymerase Mix (Takara) following manufacturers specifications.

#### Virus Purification

The wells presenting a recombinant genotyping were recovered, centrifuged 5 min at 500 × g to remove the cellular debris, and used to infect ten 75 cm² tissue-culture flasks plated with fresh *Acanthamoeba* cells. After lysis completion, the cultures were recovered, centrifuged 5 min at 500 × g to remove the cellular debris, and the virus was pelleted by a 45 min centrifugation at 6,800 × g prior purification. The viral pellet was then resuspended and washed twice in PBS and layered on a discontinuous CsCl gradient (1.2/1.3/1.4/1.5 g/cm^3^), and centrifuged at 100,000 × g overnight. An extended protocol is shown in^45^.

### Fluorescence imaging

Transfected *A. castellanii* cells were grown on poly-L-lysine coated coverslips in a 12-well plate and fixed with PBS containing 3.7% formaldehyde for 20 min at room temperature. After one wash with PBS buffer, coverslips were mounted on a glass slide with 4 μl of VECTASHIELD mounting medium with DAPI and the fluorescence was observed using a Zeiss Axio Observer Z1 inverted microscope using a 63x objective lens associated with a 1.6x Optovar for DIC, mRFP or GFP fluorescence recording.

### Statistics and reproducibility

All data are presented as the mean ± s.d. of 3 independent biological replicates (n = 3), unless otherwise stated in the figure. All data analysis were carried out using Graphpad Prism. The null hypothesis (α = 0.05) was tested using unpaired two-tailed Student’s t-tests.

### Viral fitness determination

Viral burst was calculated by manual counting using a hemocytometer chamber. In case of purified viral samples, optical density was utilized for viral quantification. Purity of the viral samples were analyzed by microscopy as previously described^45^.

### Pulsed-field gel electrophoresis

Viral suspensions were prepared according to^46^. Drops of 45 μL of the viral suspension were embedded in 1% low melting agarose, and the plugs were incubated in lysis buffer (50 mM Tris-HCl pH 8.0, 50 mM EDTA, 1% (v/v) laurylsarcosine, and 1 mg/mL proteinase K) for 24 h at 50 °C with light shaking (500 rpm). The lysis buffer was renewed every 8 h and 1 mM DTT was added 30 min before the second buffer change. After lysis, the plugs were washed once in sterile water and three times in TE buffer (10 mM Tris HCl pH 8.0 and 1 mM EDTA), for 15 min at 50 °C. Electrophoresis was carried out in 0.5× TBE using a 1% agarose gel for 20 h 10 min at 6V/cm, 120° included angle and 14 °C constant temperature in a CHEF-MAPPER system (Bio-Rad) with pulsed times ranging from 0.5 s to 3 min 10 s with a linear ramping factor.

### Quantitative PCR analysis

Viral genomes or gDNA from infected amoebas were purified using Wizard genomic DNA purification kit (PROMEGA). To determine the amplification kinetic, the fluorescence of the EvaGreen dye incorporated into the PCR product was measured at the end of each cycle using SoFast EvaGreen Supermix 2× kit (Bio-Rad, France). A standard curve using gDNA of purified viruses was performed in parallel of each experiment. For each point, a technical triplicate was performed.

### Electron microscopy

Extracellular virions or A. castellanii-infected cell cultures were fixed by adding an equal volume of PBS with 2% glutaraldehyde and 20 min incubation at room temperature. Cells were recovered and pelleted 20 min at 5,000 × g. The pellet was resuspended in 1 mL PBS with 1% glutaraldehyde, incubated at least 1 h at 4 °C, and washed twice in PBS prior coating in agarose and embedding in Epon resin. Each pellet was mixed with 2% low melting agarose and centrifuged to obtain small flanges of approximately 1mm^3^ containing the sample coated with agarose. These samples were then prepared using the osmium-thiocarbohydrazide-osmium method: 1 h fixation in 2% osmium tetroxide with 1.5% potassium ferrocyanide, 20 min in 1% thiocarbohydrazide, 30 min in 2% osmium tetroxide, overnight incubation in 1% uranyl acetate, 30 min in lead aspartate, dehydration in increasing ethanol concentrations (50, 70, 90 and 100% ethanol) and embedding in Epon-812. Ultrathin sections of 70 nm were observed using a FEI Tecnai G2 operating at 200 kV^47^.

### Comparative genomics and phylogeny

The protein-coding genes annotated in the reassembled Pandoravirus neocaledonia genome were compared to the ones of the following viral genomes: Emiliania huxleyi virus 86 (NC_007346.1), Feldmannia species virus (NC_011183.1), Ectocarpus siliculosus virus 1 (NC_002687.1), Ostreococcus tauri virus 1 (NC_013288.1), Paramecium bursaria Chlorella virus 1 (NC_000852.5), Mollivirus sibericum (NC_02 7867.1), Mollivirus Kamchatka (MN812837.1), Pandoravirus dulcis (NC_021858.1), Pandoravirus salinus (NC_022098.1), Pandoravirus quercus (NC_037667.1), Pandoravirus macleodensis (NC_037665.1) and Pandoravirus kuranda (ON887157). We used OrthoFinder (version 2.5.4) to compute the orthogroups (OG) of shared genes using the following parameters: “-M msa -S diamond_ultra_sens -y”. From a set of 438 OG selected by OrthoFinder (“Orthogroups_for_concatenated_alignment.txt” file) we kept the 435 OG without paralogs, performed a multiple alignment of each OG using Clustal omega (version 1.2.4) and concatenated the multiple alignments using Catsequence. Next a partitioned phylogeny was computed using IQ-Tree (version 2.1.2) with the “-bb 1000 -m MFP+MERGE -rcluster 10” parameters. The root of the tree was calculated using the midpoint rooting method.

### Density of shared genes along the Pandoravirus neocaledonia

Each Pandoravirus neocaledonia gene was given an “ancestry score” based on the OGs and the phylogenetic tree. This score indicates the node of the last common ancestor of all viruses having a homologue of the given gene. It ranges from 0 (genes only present in Pandoravirus neocaledonia) up to 5 (ancient, shared with at least one phycodnavirus). We next sliced the Pandoravirus neocaledonia genome in 100 bins and computed the fraction of genes of each category. Finally, these values were smoothed by computing the average values over a sliding window of 20 bins.

## Supplementary

**Figure S1.**
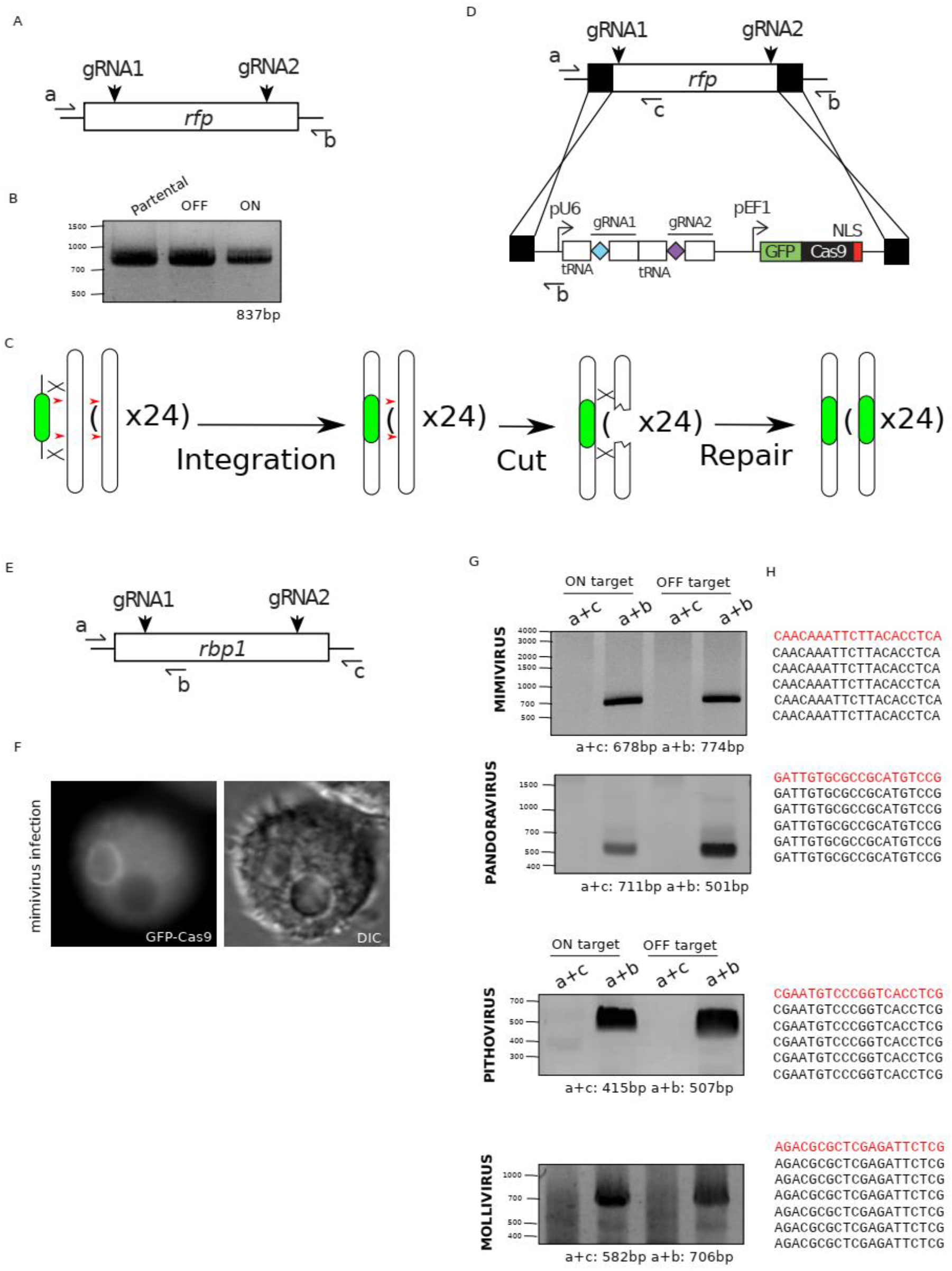
(A).Schematic representation of the rfp locus, guide targeting location and primer annealing sites. (B).PCR on the genomic locus of the rfp shows no large deletions. (C).Schematic representation of “mutagenic chain reaction” strategy. Integration of CRISPR/Cas9 in the first locus would trigger the double strand break in other alleles. Repair by homologous recombination would allow the production of homozygosis. (D).Schematic representation of the *rfp* locus, guide targeting location homology arms for recombination and primer annealing sites for a disruption using a “mutagenic chain reaction” strategy. (E).Schematic representation of the rbp1 locus, guide targeting location and primer annealing sites. (F).Representative micrograph showing exclusion of GFP-Cas9 from the viral factory of mimivirus. (G).PCR on the genomic locus of the rbp1 shows no noticeable deletions. (H).Sequencing results from a library from the pcr products demonstrate lack of mutant viruses.

**Figure S2.**
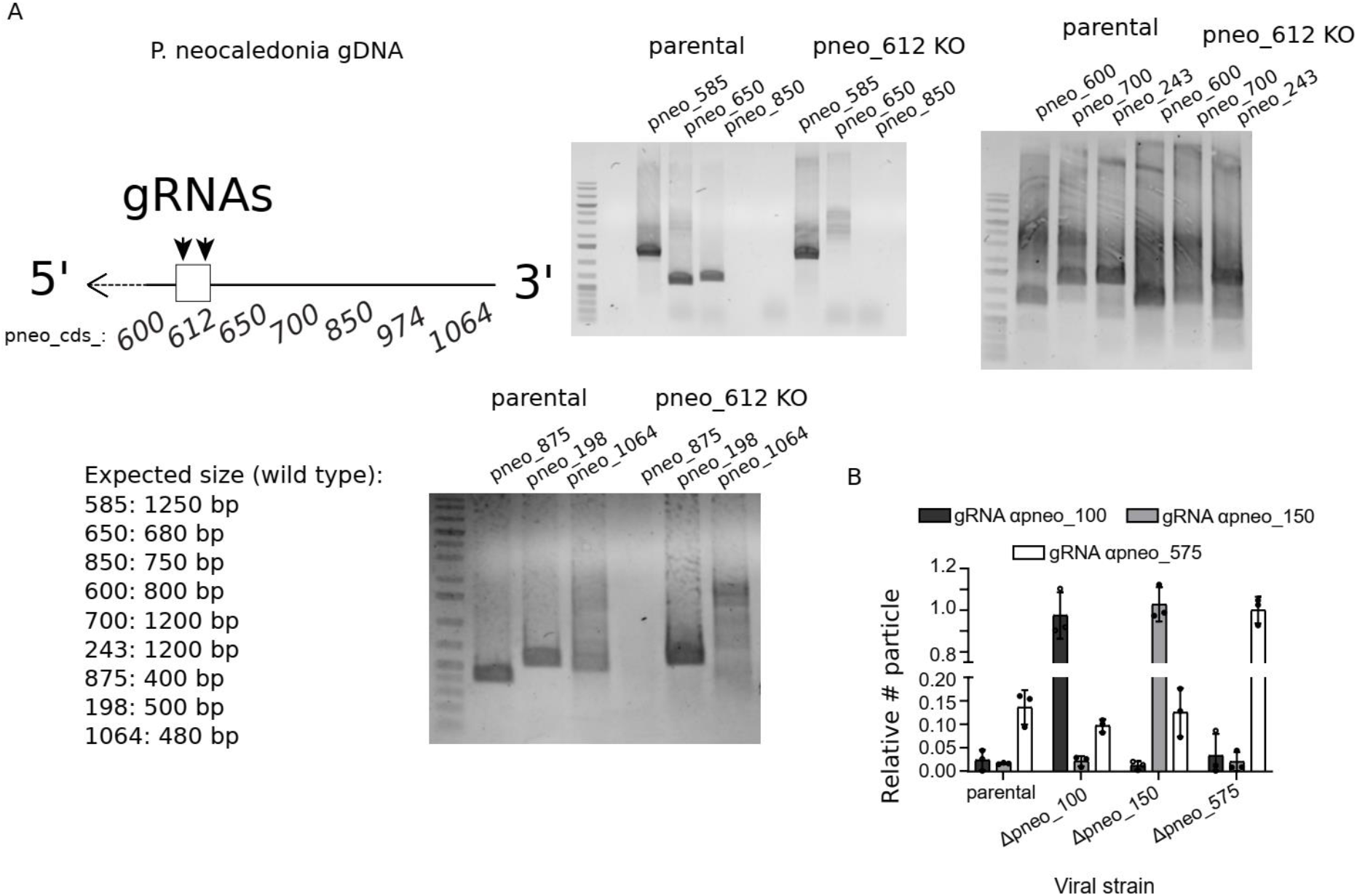
(A).PCR on different genomic locus shows large deletion at the 3’ end of pandoravirus. (B).Relative quantification of the number of viruses produced 24 hpi of CRISPR/Cas9 modified viral strains. Guides off-target and on-target of different genes were used. Data correspond to the mean ± SD of 3 independent experiments. α indicates the target of the gRNA.

**Figure S3.**
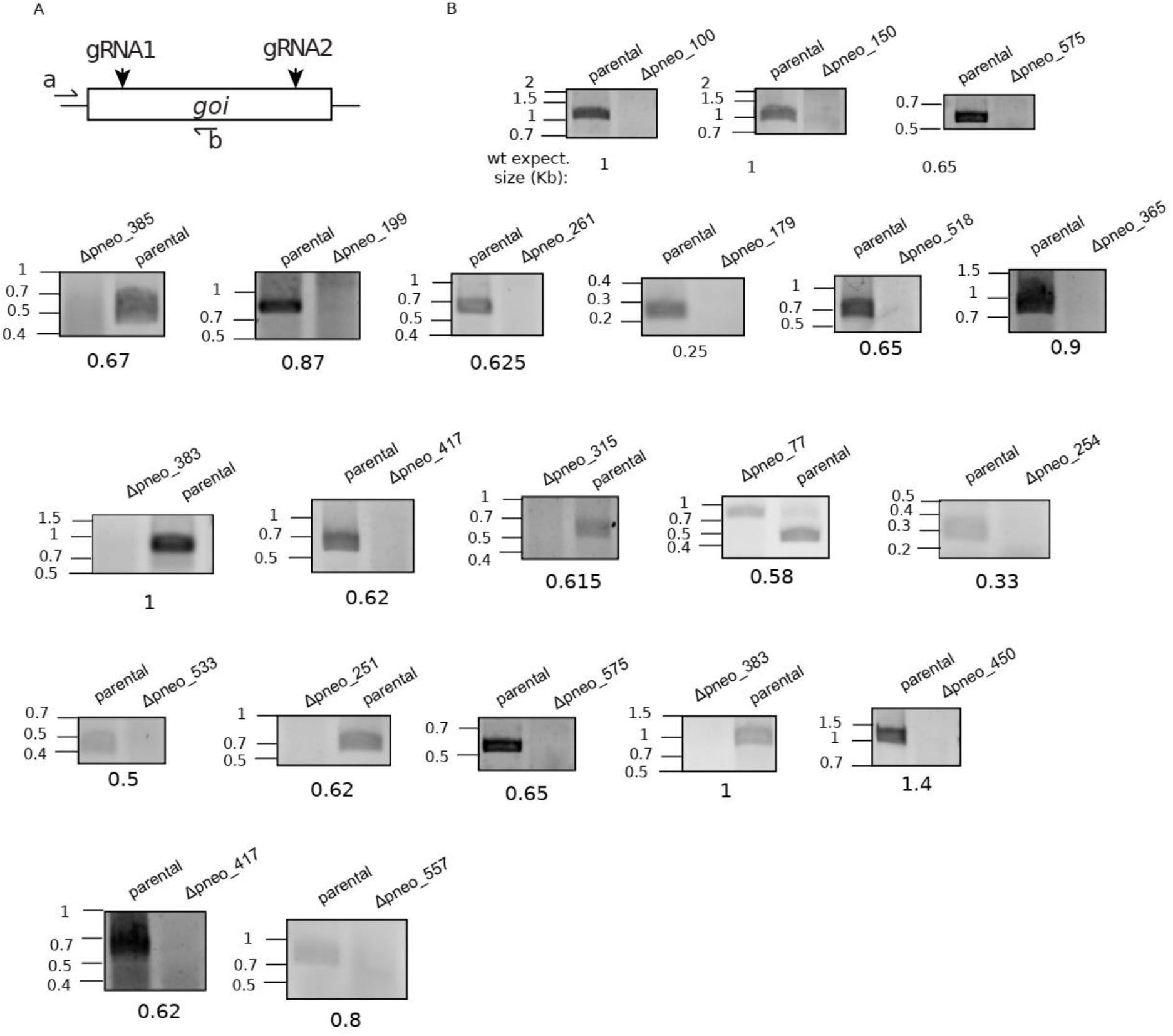
(A).Schematic representation of the *gene of interest* (*goi*) locus, guide targeting location and primer annealing sites. B.PCR on the genomic locus of the *goi*.

**Figure S4.**
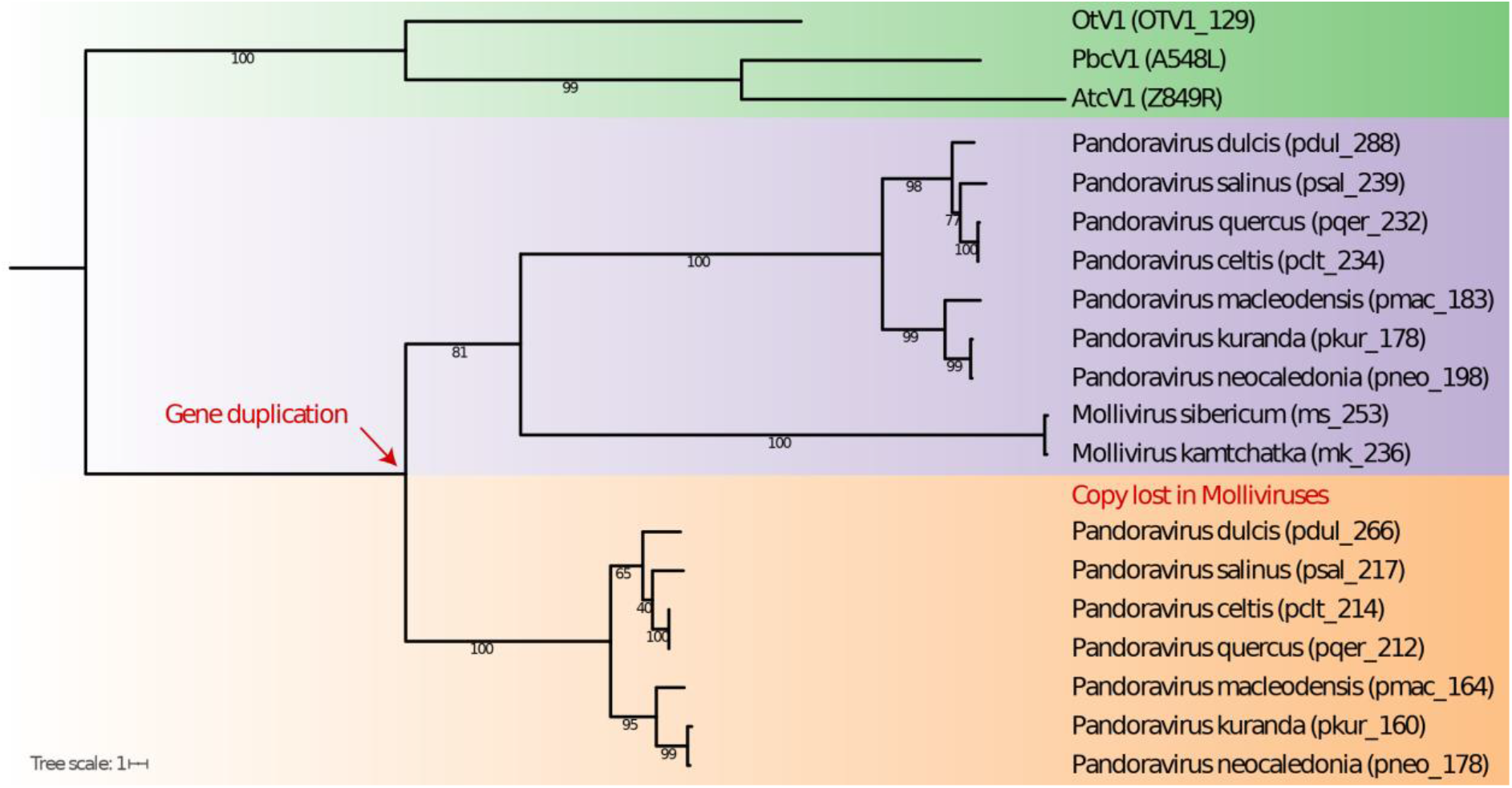
(A) Phylogenetic analysis of the SNF2-like helicases in phycodnavirus, mollivirus and pandoravirus. Emiliania huxleyi virus 86 (NC_007346.1), Feldmannia species virus (NC_011183.1), Ectocarpus siliculosus virus 1 (NC_002687.1), Ostreococcus tauri virus 1 (NC_013288.1), Paramecium bursaria Chlorella virus 1 (NC_000852.5), Mollivirus sibericum (NC_02 7867.1), Mollivirus Kamchatka (MN812837.1), Pandoravirus dulcis (NC_021858.1), Pandoravirus salinus (NC_022098.1), Pandoravirus quercus (NC_037667.1), Pandoravirus macleodensis (NC_037665.1) and Pandoravirus kuranda (ON887157).

**Figure S5.**
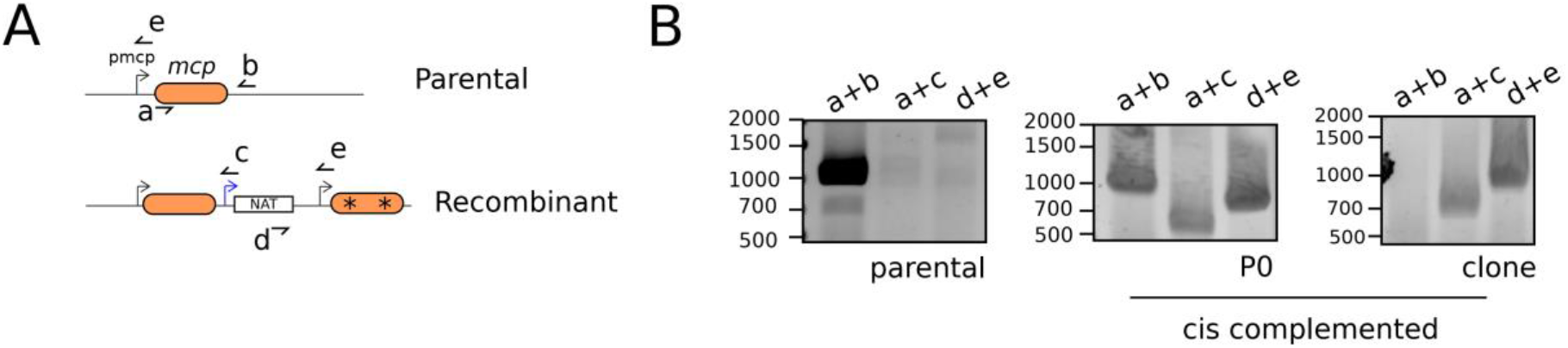
(A).Schematic representation of the *mcp* locus and primer annealing sites. B.PCR on the genomic locus of the *mcp* demonstrate correct integration and clonality. Integration in the population at the generation 0 (P0, 24hours posttransfection) is also shown. Expected size for pcr products: a+b (parental): 1000bp, a+c (recombinant): 650bp, d+e (recombinant): 800bp.

**Table S1:List of targeted genes on Pandoravirus neocaledonia classified according to their broad function.**
NA: non-assessed, D: Dispensable and E: essential.

**Table S2:List of primers used in this study.**

**Table S3:List of constructs generated in this study.**

